# A Model for DHX15 Mediated Disassembly of A-Complex Spliceosomes

**DOI:** 10.1101/2021.09.10.459862

**Authors:** Hannah M. Maul-Newby, Angela N. Amorello, Turvi Sharma, John H. Kim, Matthew S. Modena, Beth Prichard, Melissa S. Jurica

**Author notes:** Corresponding author (MSJ).

## Abstract

A critical step of pre-mRNA splicing is the recruitment of U2 snRNP to the branch point sequence of an intron. U2 snRNP conformation changes extensively during branch helix formation and several RNA-dependent-ATPases are implicated in the process. However, the molecular mechanisms involved remain to be fully dissected. We took advantage of the differential nucleotide triphosphates requirements for DExD/H-box enzymes to probe their contributions to in vitro spliceosome assembly. Both ATP and GTP hydrolysis support the formation of A-complex, indicating the activity of a DEAH-enzyme because DEAD-enzymes are selective for ATP. We immunodepleted DHX15 to assess its involvement and although splicing efficiency decreases with reduced DHX15, A-complex accumulation incongruently increases. DHX15 depletion also results in the persistence of the atypical ATP-independent interaction between U2 snRNP and a minimal substrate that is otherwise destabilized in the presence of either ATP or GTP. These results lead us to hypothesize that DHX15 plays a quality control function in U2 snRNP’s engagement with an intron. In efforts to identify the RNA target of DHX15, we determined that an extended polypyrimidine tract is not necessary for disruption of the atypical interaction between U2 snRNP and the minimal substrate. We also examined U2 snRNA by RNase H digestion and identified nucleotides in the branch binding region that become accessible with both ATP and GTP hydrolysis, again implicating a DEAH-enzyme. Together, our results demonstrate that multiple ATP-dependent rearrangements are likely involved in U2 snRNP addition to the spliceosome and that DHX15 can have an expanded role in splicing.

## Introduction

Pre-mRNA splicing by the spliceosome is an essential step in eukaryotic gene expression and must be highly accurate to generate functional messenger RNAs. The boundaries of an intron are initially designated by base-pairing interactions between U1 snRNA and the 5’ splice site and U2 snRNA and the branch point sequence. U2 snRNA recognition of the branch point sequence takes place in the context of a small ribonucleoprotein particle (snRNP), which also contains ten core proteins, three SF3A proteins, and seven SF3B proteins. These events signal the rest of spliceosome to assemble into a catalytic entity.

A collection of RNA-dependent ATPases that drive rearrangements required during spliceosome assembly have also been linked to enforcing splice site fidelity in *S. cerevisiae* (Koodathingal et al. 2010; Koodathingal and Staley 2013; Mayas et al. 2006; Toroney et al. 2019; Villa and Guthrie 2005; Xu and Query 2007; Yang et al. 2013). Most of these enzymes, classified as either DEAD (DDX) or DEAH (DHX), are considered helicases because they can disrupt RNA base-pairing interactions, however they exhibit some mechanistic differences (Gilman et al. 2017; Jankowsky 2011). DDX enzymes have a Q-motif that enforces a selectivity for ATP as the source of energy for RNA unwinding. In contrast, DHX enzymes lack this motif and can utilize other nucleotides, with a preference for ATP and GTP.

During *in vitro* spliceosome assembly, recruitment of U2 snRNP to the branch point sequence is the first ATP-dependent step, and results in formation of a stable entity referred to as A-complex. The mechanistic basis of the requirement for ATP is not fully understood, but striking differences between recent cryo-EM structures of U2 snRNP and A-complex imply several large-scale molecular rearrangements of interactions within the snRNP and with the intron. Several RNA-dependent ATPases are associated with U2 snRNP: three DDX-enzymes (DDX46, DDX39B, DDX42) and DHX15. Both DDX46 (aka Prp5) and DDX39B (aka UAP56 or Sub2) have been linked to A-complex assembly, and likely mediate some of the rearrangements. For example DDX46 has been localized near the branch stem loop (BSL) in the 17S U2 snRNP structure (Zhang et al. 2020), and in yeast BSL mutations suppress an N-terminal deletion of the DDX46 ortholog Prp5 (Perriman and Ares 2010). DDX46 is also proposed to proofread the extended helix that forms between U2 snRNA and the branch point sequence of an intron (Liang and Cheng 2015; Xu and Query 2007; Zhang et al. 2021). UAP56 interacts with U2AF, a factor that bind the polypyrimidine tract (PYT) downstream of the branch to help recruit U2 snRNP (Fleckner et al. 1997). DDX42 associates exclusively with a form of U2 snRNP that lacks the SF3A and SF3B complexes and is unlikely to be directly involved in spliceosome assembly (Will et al. 2002). DHX15 (aka Prp43) is best known for its role in disassembly of the intron lariat spliceosome (ILS) at the end of splicing, although studies differ on whether it interacts directly with the intron or U6 snRNA (Fourmann et al. 2016; Toroney et al. 2019). DHX15 also has a role in ribosome biogenesis, and its specificity appears to be regulated by different G-patch cofactors that direct it to the intron lariat (Wen et al. 2008) or pre-rRNA (Memet et al. 2017). Notably, DHX15 is present in pulldowns of U2 snRNP and A-complex spliceosomes, along with G-patch proteins RBM5, RBM10, RBM17, CHERP and SUGP1 (Agafonov et al. 2011).

For human introns, the sequences that define splice sites are quite variable, especially the branch point sequence. Quality control of branch point selection is likely important to maintain cellular health as alterations in the players involved is evidenced to lead to cancer (Bonnal et al. 2020). Unsurprisingly, these include mutations in several proteins associated with U2 snRNP. For example, specific point mutations in SF3B1, a U2 snRNP protein that directly contacts the branch point sequence in early spliceosome assembly, are enriched in cells from cancers and dysplasia syndromes, particularly in hematological lineages (Dvinge et al. 2016). In addition, SF3B1 mutations result in altered branch point selection for some introns (Alsafadi et al. 2016; Darman et al. 2015; DeBoever et al. 2015; Kesarwani et al. 2017; Wang et al. 2016).

In this study, we use an *in vitro* splicing system to investigate the possibility of a role for DHX15 in mediating ATP-dependent rearrangements involved in U2 snRNP’s engagement with an intron. Depletion of DHX15 results in increased A-complex formation, but lower splicing efficiency. The effect is amplified when we assemble spliceosomes on introns incapable of forming complexes competent to complete splicing, leading us to propose that DHX15 may have an additional splicing related function in quality control of branch sequence recognition. Employing a powerful affinity-tagged version of the core component SNRPB2, we also characterize changes in U2 snRNP that show differential selectivity for ATP and GTP, suggesting involvement of different DDX and DHX enzymes in a growing constellation of rearrangements that may impact how the spliceosome finds introns with high fidelity.

## Results

### Both ATP & GTP can support spliceosome assembly and splicing

To test the hypothesis that molecular rearrangements during A-complex assembly are promoted by one or more RNA-dependent ATPases associated with U2 snRNP, we used nucleotide preference to differentiate between the activity of DDX and DHX enzymes. DDX enzymes require ATP to disrupt RNA interactions and are often locked onto their target when bound to a non-hydrolysable ATP analog (Gilman et al. 2017). In contrast, DHX enzymes are processive RNA helicases that can also utilize GTP (Jankowsky 2011). If a DHX enzyme is involved in the initial rearrangement, then GTP should support A-complex formation prior to the first rearrangement that requires DDX enzyme activity. Alternatively, if ATP-binding is sufficient for an initial DDX-mediated rearrangement, then non-hydrolysable AMP-PNP will support assembly to that point.

We carried out *in vitro* splicing assays on a radiolabeled pre-mRNA substrate in the presence or absence of ATP, GTP and AMP-PNP and excluded creatine phosphate to prevent recycling of nucleotides. We used native gels to assess spliceosome assembly on the pre-mRNA and denaturing gels to measure splicing efficiency (Fig. 1A and B). As expected, ATP confers robust spliceosome assembly and splicing chemistry, which are both lost with no added nucleotide. AMP-PNP does not support assembly nor splicing, leading us to conclude that binding of a target RNA by a DDX enzyme is not sufficient to promote the first rearrangement needed for U2 snRNP to engage an intron. Surprisingly, GTP promotes both a significant amount of spliceosome assembly and splicing, although less efficiently than with ATP. We also tested nucleotide requirements in the context of a truncated substrate (A^3’^ substrate) that recapitulates U2 snRNP interactions with an intron (Konarska and Sharp 1986). We find that both ATP and GTP addition results in formation of the A^3’^-complex, but not AMP-PNP (Fig. 1C). Because GTP supports A-complex assembly, we conclude that a DHX enzyme can promote an initial rearrangement that allows U2 snRNP to engage with an intron. We hypothesize that the higher accumulation of A-complex with ATP reflects the contribution of a DDX enzyme, potentially by capturing a transient rearrangement or conformation to further promote spliceosome assembly. Based on their direct association with U2 snRNP, DHX15 and DDX46 are most-likely the candidates for the two activities.

**Figure 1:**
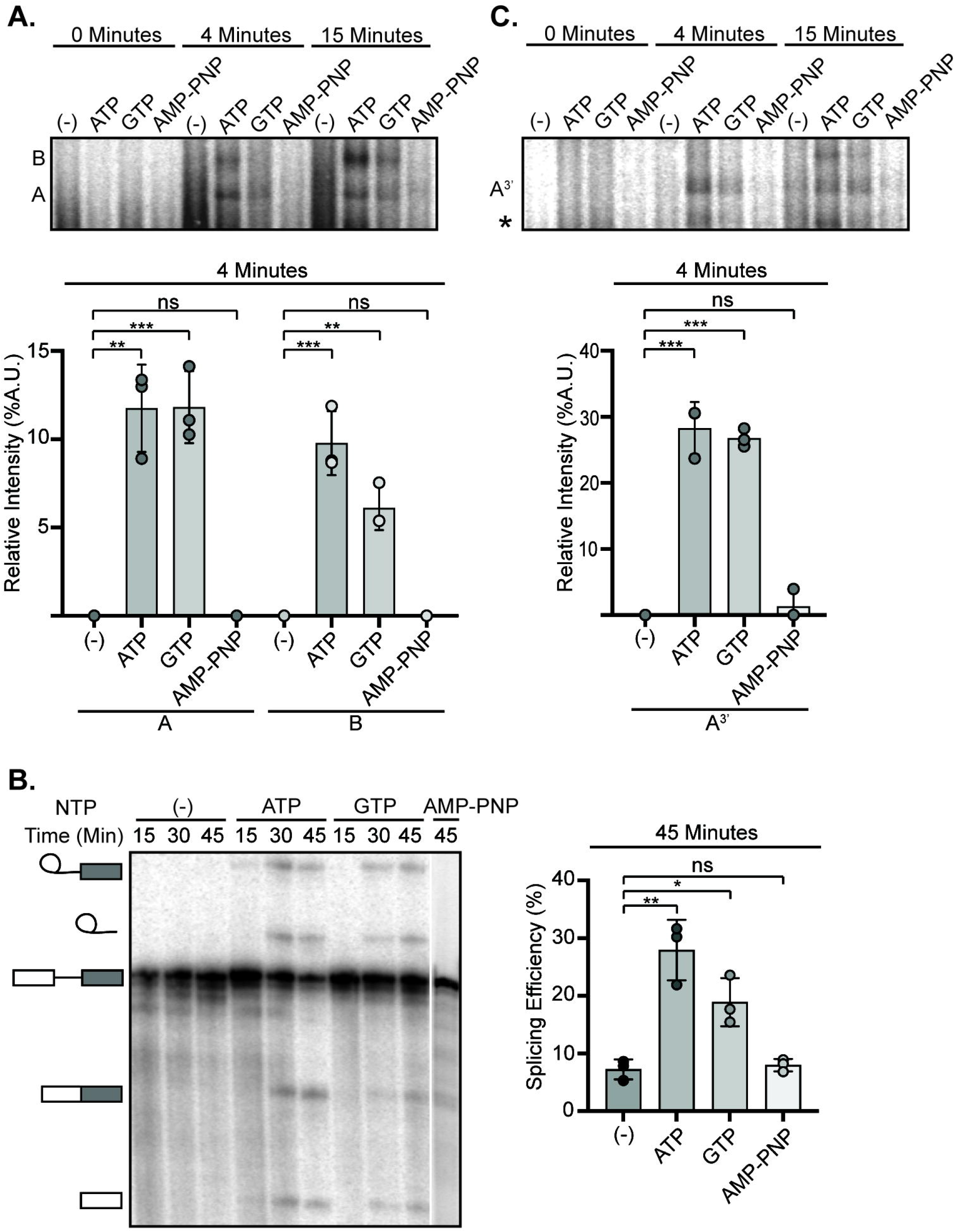
Both ATP & GTP can support spliceosome assembly and splicing. A. Top: Representative native gel analysis of *in vitro* spliceosome assembly with a radiolabeled full-length substrate with different added NTPs at the indicated timepoints. A- and B-complex bands positions are labeled. Bottom: Normalized band intensity relative to the entire lane is shown for the 4-minute timepoint for three independent trials. Statistical differences were examined by unpaired Student’s t-test with *** p<0.001, ** p<0.01, *p<0.05. B. Same as (A) except that an A^3’^-substrate was used for spliceosome assembly. C. Left: Representative denaturing gel analysis of radiolabeled RNA isolated at the indicated timepoints from *in vitro* splicing reactions using a full-length substrate with different added NTPs. The RNA band identities are illustrated on the left as (top to bottom): lariat intron intermediate, lariat intron, pre-mRNA substrate, mRNA, and 5’ exon intermediate. Right: Splicing efficiency is measured as intensity of the mRNA band over total RNA bands and shown for the 45-minute timepoint for three independent trials. Statistical differences were examined as in (A).

### Depletion of DHX15 reduces splicing efficiency, but not spliceosome assembly

To determine whether DHX15 has a role in U2 snRNP addition to the spliceosome, we immunodepleted the protein from HeLa nuclear extract. Despite the high abundance of DHX15, 55-72% of the protein was removed in three independent immunodepletions relative to mock depletion as determined by western blot analysis (Fig. 2A, Sup. Fig. 1). We used DHX15- and mock-depleted extracts for *in vitro* spliceosome assembly on the A^3’^ substrate to focus on U2 snRNP incorporation. Contrary to the prediction that loss of DHX15 would inhibit assembly, the relative band intensity of A-complex in DHX15-depeleted extract is increased compared to the mock-depleted extract (Fig. 2B). We repeated the experiment with a full-length substrate to determine if the difference is due to the truncated substrate. A small but consistent increase in the relative band intensities of both A- and B-complex spliceosomes persist with DHX15-depleted extracts (Fig. 2C). Surprisingly, overall splicing efficiency is lower in the DHX15-depleted extracts compared to mock-depleted extract (Fig. 2D).

**Figure 2:**
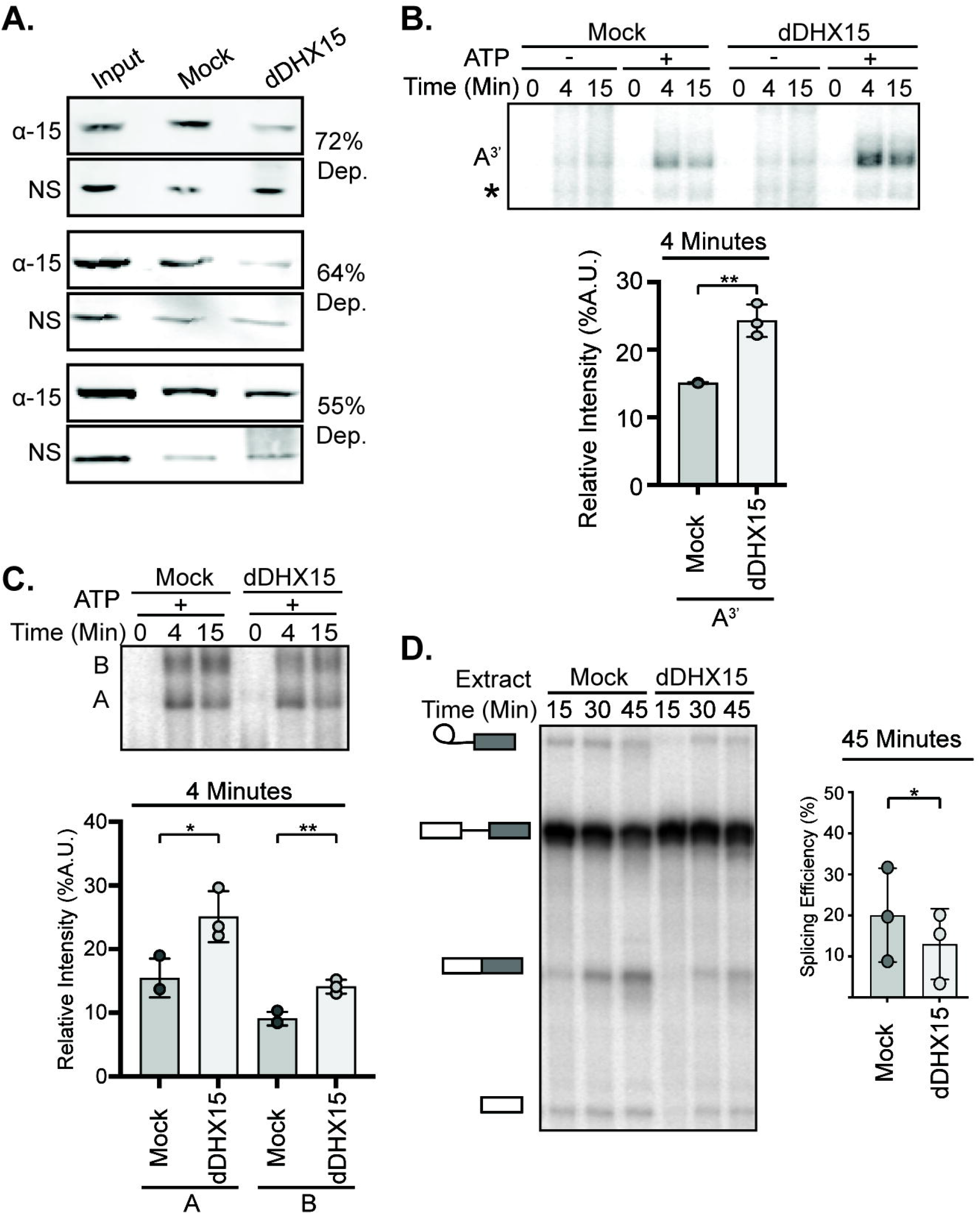
Depletion of DHX15 reduces splicing efficiency, but not spliceosome assembly. A. Western blot analysis of three independent immunodepletions of DHX15 from HeLa nuclear extract. Depletions were quantified by comparing the band intensity of DHX15 (α-15) to a non-specific band (NS), both normalized to the intensity of the entire lane. Samples are shown for untreated nuclear extract (Input), control depletion with 1X PBS or IgG (Mock) and depletion with anti-DHX15 (dDHX15). B. Top: Representative native gel analysis of *in vitro* spliceosome assembly with a radiolabeled A^3’^-substrate in Mock and dDHX15 nuclear extracts at the indicated timepoints. Bottom: Normalized band intensity relative to the entire lane is shown for the 4-minute timepoint from three independent trials. Statistical differences were examined by paired Student’s t-test with p-values as in Figure 1A. C. Same as (B) but with a full-length splicing substrate. D. Left: Representative denaturing gel analysis of radiolabeled RNA isolated at the indicated timepoints from *in vitro* splicing reactions using Mock or dDHX15 nuclear extracts. The RNA band identities are as described in Fig 1C. Right: Splicing efficiency determined as in Fig. 1C.

In the context of our original hypothesis, these results indicate that DHX15 may not be responsible for the initial rearrangement that allows U2 snRNP to interact with an intron, although we cannot rule out that the remaining DHX15 after immunodepletion is sufficient to carry out that role. However, the opposing increase in spliceosome assembly and decreased splicing efficiency in DHX15-depleted extracts lead us to suspect that DHX15 may have a role in quality control of early spliceosome assembly. We postulate that DHX15 promotes disassembly of unproductive spliceosome complexes. With reduced DHX15 activity, these unproductive spliceosomes accumulate and potentially sequester spliceosome components, which results in overall less splicing efficiency. Alternatively, the decrease in splicing efficiency may indirectly result from the loss of spliceosome disassembly in the ILS or yet another role for DHX15 after B-complex formation that promotes splicing chemistry. As we are not able to add back purified DHX15, we also cannot rule out the possibility that a co-depleted factor is responsible for the observed effects.

### Reduction of DHX15 stabilizes the ATP-independent interaction between U2 snRNP and minimal intron

Query *et al*. showed a minimal RNA (A^min^ substrate) containing only a branch point sequence followed by a PYT interacts with U2 snRNP in the absence of ATP to form the A^min^-complex (Query et al. 1997). Notably, in the presence of ATP, the A^min^-complex is destabilized by an unknown entity (Newnham and Query 2001). To determine whether DHX15 could be responsible, we incubated the A^min^ substrate in both DHX15-depleted and mock-depleted HeLa nuclear extract with and without ATP. In the absence of ATP, the expected A^min^-complex band forms in both extracts (Fig. 3A). Addition of ATP to the mock-depleted extract results in the loss of most A^min^-complex and the appearance of a faster migrating complex of unknown composition (*) that decreases over time (Query et al. 1997). In DHX15-depleted extracts with ATP, the A^min^-complex persists, indicating that DHX15 and/or a co-depleted factor is responsible for the ATP-dependent loss. The faster migrating complex (*) is largely unaffected.

**Figure 3:**
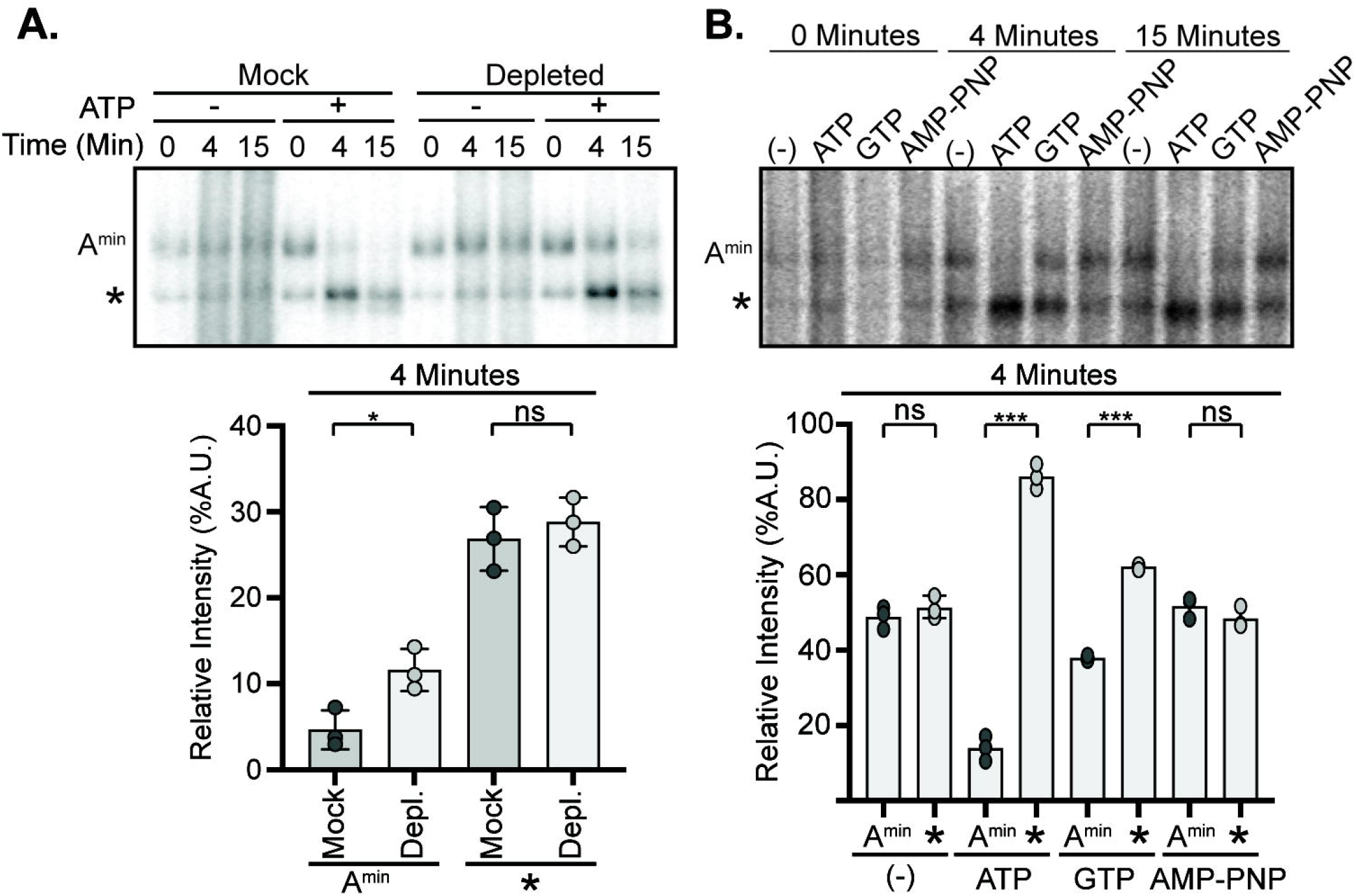
Reduction of DHX15 stabilizes the ATP-independent interaction between U2 snRNP and minimal intron. A. Top: Representative native gel analysis of *in vitro* spliceosome assembly with a radiolabeled A^min^-substrate in control (Mock) and DHX15-immunodepleted (dDHX15) nuclear extracts at the indicated timepoints. Bottom: Normalized band intensity relative to the entire lane is shown for the 4-minute timepoint from three independent trials. Statistical differences were examined by paired Student’s t-test. Top: Representative native gel analysis of *in vitro* spliceosome assembly with A^min^-substrate with different added NTPs at the indicated timepoints. Statistical differences were examined by unpaired Student’s t-test.

If DHX15 is responsible for disrupting the unproductive A^min^-complex, then GTP should also promote its disassembly in normal nuclear extract. We compared the addition of ATP, GTP and AMP-PNP with the A^min^ substrate, and found that both ATP and GTP result in the loss of A^min^-complex, while AMP-PNP does not (Fig. 3B). Taken together, these results support a model in which DHX15 disrupts an unproductive interaction between U2 snRNP and the intron in A^min^-complex as a quality control mechanism.

### A^min^ destabilization does not occur via an extended PYT

In the A^min^-complex, there are only two possible RNAs for DHX15 to target: U2 snRNA and the short A^min^ substrate. Given that DHX enzymes act by translocating from 3’ to 5’ on a single RNA strand (Hamann et al. 2019; Mallam et al. 2012; Sengoku et al. 2006; Tauchert et al. 2017; Yang et al. 2007; Yang and Jankowsky 2006), the 3’ end of the A^min^ substrate is a likely DHX15 target. We tested this possibility by truncating the 3’ end of the intron in 4 nucleotide increments to see if loss of the putative target sequence results in A^min^-complex stabilization despite added ATP (Fig. 4A). We stopped at 10 nucleotides downstream of the branch point sequence because too short of a PYT interferes with spliceosome assembly (Bessonov et al. 2010). The truncations did not interfere with A^min^ assembly in the absence of ATP (Fig. 4B), and complexes were all destabilized with added ATP. We conclude an extended PYT is not necessary for A^min^ destabilization, meaning that DHX15 either interacts with the remaining 10 nucleotides of the PYT or recognizes another RNA feature within the A^min^-complex.

**Figure 4:**
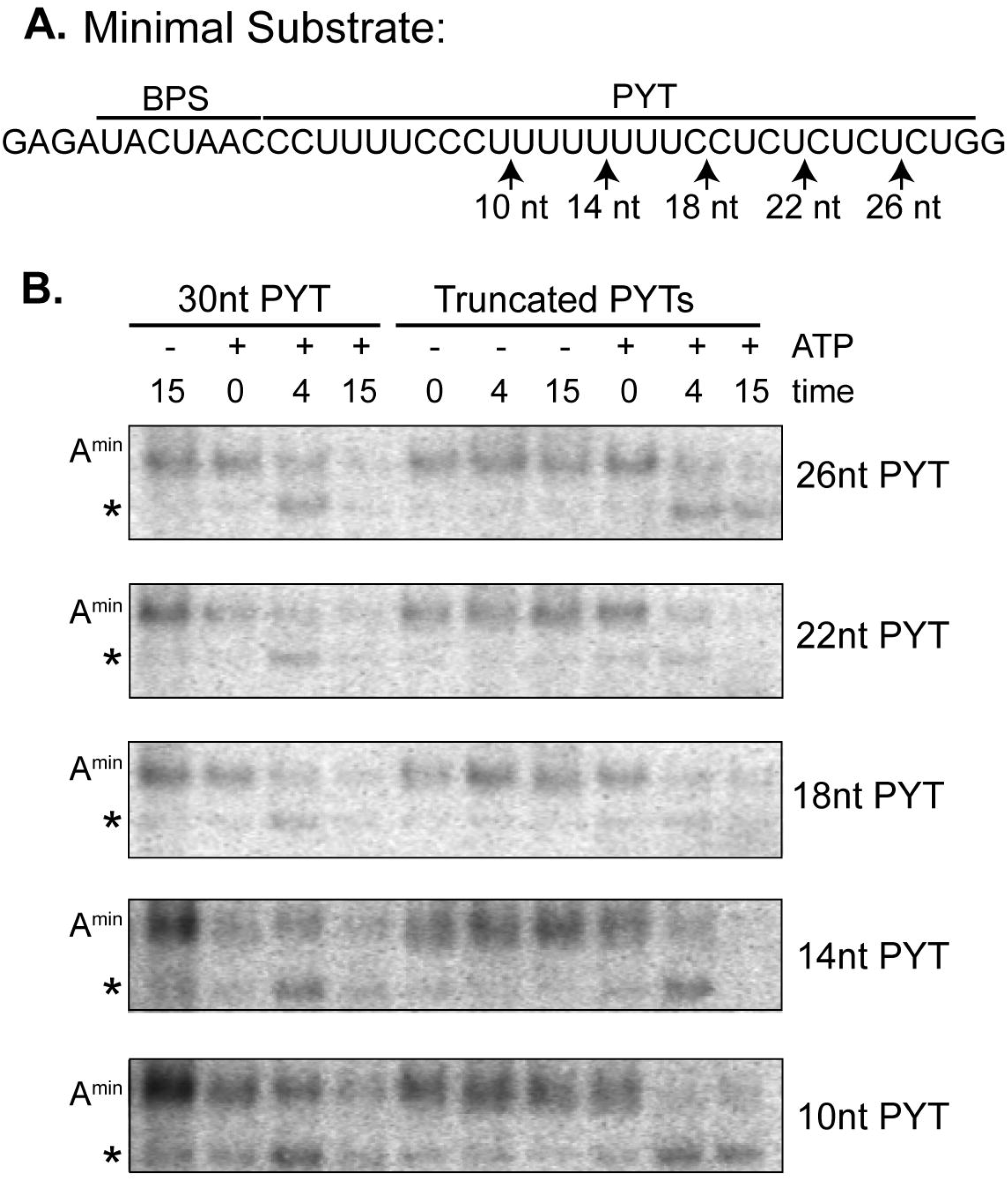
Amin destabilization does not occur via an extended PYT. A. Schematic of the A^min^ RNA substrate with branch point sequence (BPS) and PYT. The arrows indicate the 3’ end of the PYT in each RNA. B. Top: Representative native gel analysis of *in vitro* spliceosome assembly with truncated A^min^ substrate +/-ATP at the indicated timepoints. PYT length is indicated on the right.

### U2 snRNA accessibility is regulated by NTP hydrolysis

The other RNA in A^min^ is U2 snRNA, of which nucleotides 32-46 are of special interest because they both interact with the intron in an extended branch helix and form the mutually exclusive branch stem loop (BSL) structure (Perriman and Ares 2010; Zhang et al. 2020) (Fig. 5A). In the context of the U2 snRNP in nuclear extract, this region was shown to become accessible for base-pairing with a complementary DNA oligonucleotide in an ATP-dependent manner (Black et al. 1985), although the factor responsible for the ATP-dependence was unknown. To determine whether a DHX enzyme could be involved, we tested whether GTP or AMP-PNP could also enable base-pairing of different regions in the 5’ half of U2 snRNA. We created a series of overlapping DNA oligonucleotides complementary to U2 snRNA (nt 1-15, nt 12-26, nt 24-38, nt 32-46, Fig. 5A), which we added to HeLa nuclear extract treated with various NTP’s. If the targeted region is accessible for base-pairing, endogenous RNase H in the extracts will induce cleavage of the RNA/DNA hybrid. We mapped the specific cleavage sites using primer extension (Fig 5B) and quantified overall digestion of U2 snRNA induced by the oligonucleotides (Fig 5C).

**Figure 5:**
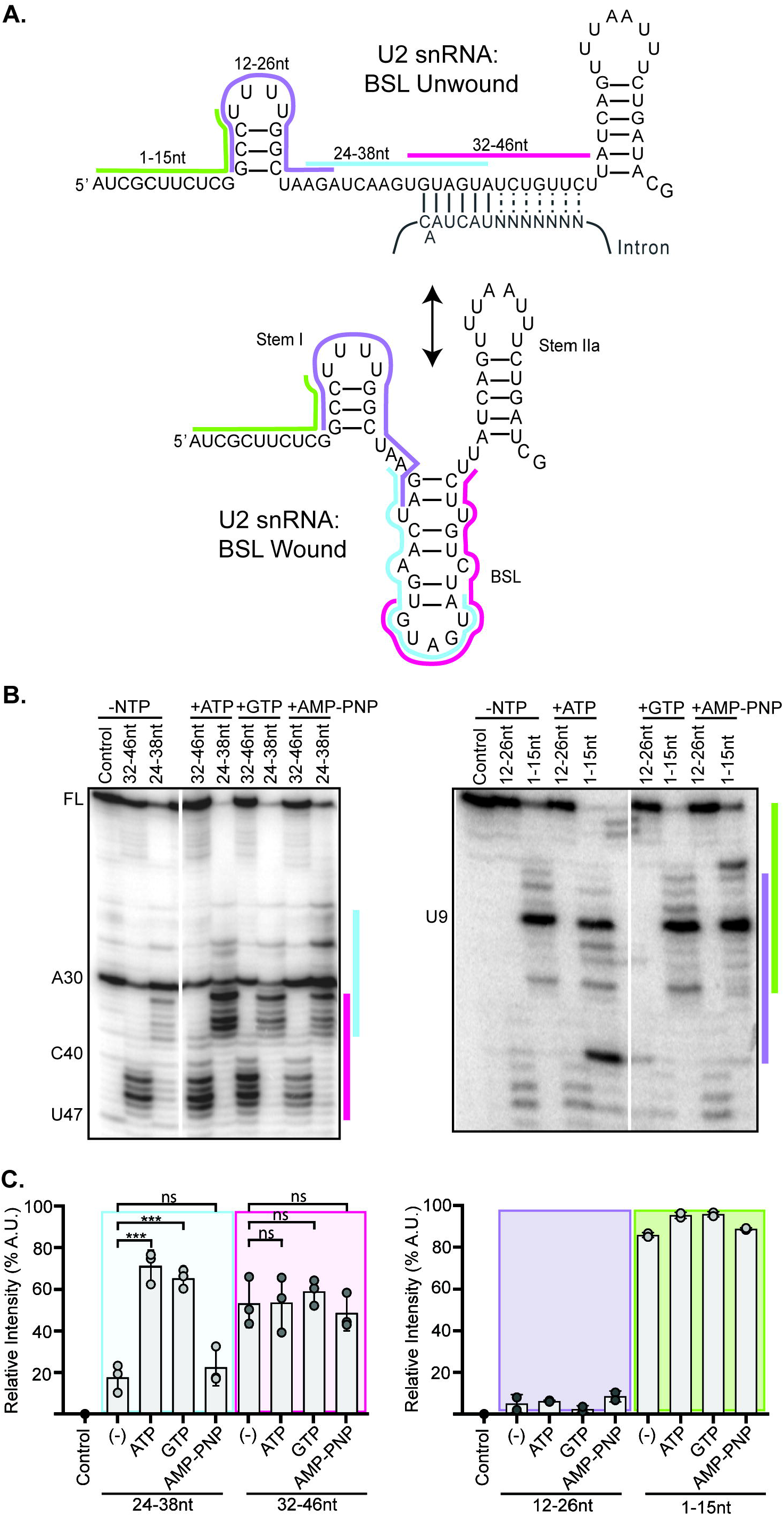
U2 snRNA accessibility is regulated by NTP hydrolysis. A. Secondary structure of the U2 snRNA with the BSL (bottom) or unwound interaction with the grey intron (top). The position of complementary DNA oligonucleotides used for RNase H digestion are indicated with colored bars. B. Primer extension analysis U2 snRNA isolated from nuclear extract after RNase H digestion. Extracts were treated with ATP, GTP or AMP-PNP or no NTP. Control digestions were carried out with non-complementary DNA oligonucleotide. C. Cleavage efficiency was determined as band intensity in the region complementary to the oligonucleotide over the intensity of the entire lane of three independent trials. Statistical differences relative to the –NTP sample were examined by unpaired Student’s t-test.

With an oligonucleotide that targets the 5’ half of the BSL (nt 24-28), over 60% of U2 snRNA molecules in the extract are cleaved after both ATP and GTP treatment, compared with ∼20% cleaved with no added NTP or AMP-PNP. Specifically, most cleavage occurs after nucleotides 32-36 (GUGUA) with some cleavage after nucleotide 37. Using an oligonucleotide targeting the 3’ half of the BSL (nt 32-46), around half of U2 snRNA molecules are cleaved, primarily after nucleotides 42-46 independent of added NTP, although there is a small increase in cleavage after nucleotides 37 and 38 with ATP and GTP. Cleavage in the presence of the other oligonucleotides is not influenced by NTP treatment. With an oligonucleotide targeting the beginning of U2 snRNA (nt 1-15), nearly all the U2 snRNA molecules are cleaved after U9 indicating that while the 5’ end of U2 snRNA is available for base-pairing, nucleotides 10-15 are protected, likely because of their participation in Stem I (Fig. 5A). An oligonucleotide that targets nucleotides 12-26 also does not induce cleavage, further supporting the stability of Stem I.

Because both ATP and GTP hydrolysis results in increased accessibility, we conclude that a DHX enzyme mediates the rearrangement that makes U2 snRNA nucleotides 24-38 available for base-pairing. The enzyme may unwind an RNA structure, such as the BSL, or dislodge a protein that protects the RNA.

### U2 snRNP composition is affected by NTP hydrolysis

To determine if a change in protein interactions is responsible for the NTP-dependent change in U2 snRNA accessibility, we generated a HeLa cell line with a stably integrated transgene encoding a V5-tagged SNRPB2, a core U2 snRNP protein, for pulldown studies (Khandelia et al. 2011; Kim 2019). Western analysis of nuclear extract shows that over half the SNRPB2 expressed in these cells carries the tag (Fig.6A). We analyzed anti-V5 immunoprecipitations (IP’s) for RNA and see enrichment of U2 snRNA indicating that the tagged protein incorporates into U2 snRNP (Fig 6B). IP’s of *in vitro* splicing reactions using a radiolabeled full-length pre-mRNA reveal higher levels of pre-mRNA co-eluting with the V5-tagged SNRPB2 relative to IgG control, which is further enhanced by addition of ATP to the splicing reactions (Fig. 6C). We repeated the IP’s with the A^min^ substrate, and in this case more RNA co-elutes from splicing reactions without added ATP (Fig. 6D). These results parallel our observations by native gel analysis, indicating that the V5-tagged U2 snRNP incorporates splicing complexes.

**Figure 6:**
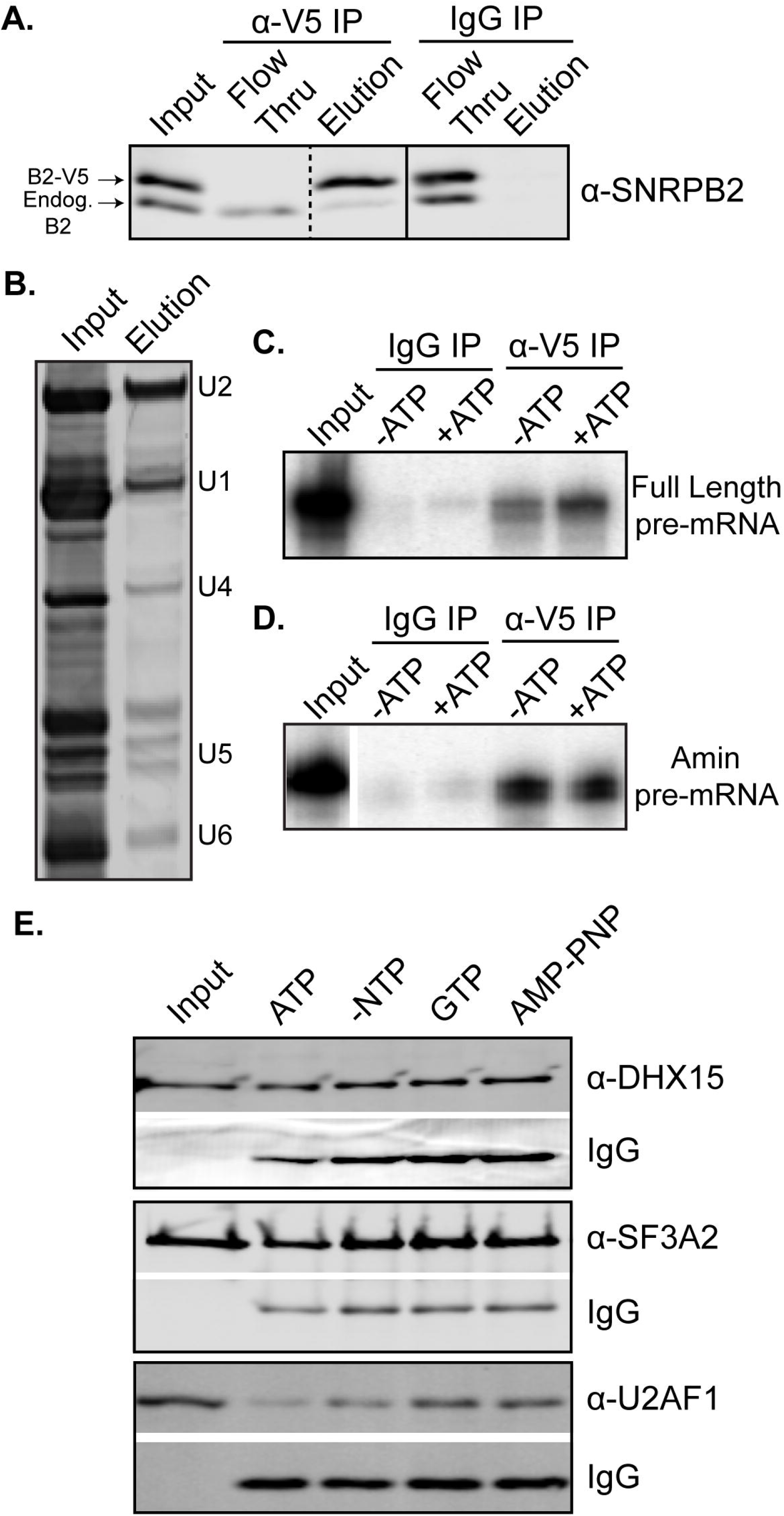
ATP hydrolysis alters U2 snRNP protein composition. A. Western blot analysis of IP with anti-V5 and mouse IgG from HeLa nuclear extract containing V5-tagged SNRPB2 and probed with anti-SNRPB2, which detects both endogenous (endog) and V5-tagged SNRPB2 (V5-B2). B. Denaturing gel analysis of RNA isolated from anti-V5 IP eluate stained with SYBR Gold. C. Denaturing gel analysis of RNA isolated anti-V5 IP of *in vitro* splicing reactions +/-ATP using a radiolabeled full-length RNA substrate in the conditions. D. Same a (C) except with an A^min^ substrate E. Western analysis anti-V5 of IP’s from nuclear extract incubated under *in vitro* splicing conditions in the presence of ATP, GTP or AMP-PNP. Blots were probed with the indicated antibody.

To test whether protein composition of U2 snRNP is altered by NTP hydrolysis, we IP’d V5-tagged U2 snRNP from nuclear extract incubated with ATP, GTP or AMP-PNP. Western analysis of eluates showed that the NTP treatments do not alter the immunoprecipitation of V5-tagged SNRPB2, or the association of the 17S U2 snRNP protein SF3A2 (Fig. 6E). We detect DHX15 in the eluates under all conditions. In contrast, U2AF1 is reduced after ATP treatment. This change is more likely due to the activity of a DDX enzyme because GTP treatment does not result in the same decrease. AMP-PNP also has no effect. Relating these results to the increased U2 snRNA accessibility in the snRNP after both ATP and GTP treatment, the ATP-specificity of U2AF1 loss means it is not responsible for blocking access to the branch binding region. Because DHX15 remains with the snRNP, it could be responsible removing a different protein to make the branch binding region of U2 snRNA available for base-pairing interactions. It is tempting to speculate that DHX15 may be disrupting unproductive spliceosomes in a similar manner.

## Discussion

Spliceosome assembly relies on an assortment of DDX and DHX enzymes to both drive molecular rearrangements and promote splicing fidelity. Both DDX46 and DDX39B play roles in U2 snRNP’s addition to the branch point sequence, although what they target, what they rearrange, and how they are regulated is still not clear. Using the differential nucleotide triphosphate selectivity of DDX- and DHX-enzymes, our study supports added roles for a DHX-enzyme(s) in regulating U2 snRNP structure and early assembly of human spliceosomes. We show that GTP can substitute for ATP in overcoming whatever blocks U2 snRNP from binding an intron with an anchor sequence. GTP can also mediate remodeling of the U2 snRNP to expose the branch binding sequence. Finally, GTP also promotes the destabilization of the unproductive interaction between U2 snRNP and an anchorless minimal intron, which we linked to the presence of DHX15. Additionally, we identified an ATP-specific loss of U2AF1 from U2 snRNP, which suggests the activity of a DDX-enzyme. DDX39B is a strong candidate because it has been shown to directly interact with U2AF1 (Shen et al. 2007).

DHX15 is a ubiquitous nuclear RNA-dependent NTPase in disparate cellular functions and is best characterized as promoting ribosomal RNA biogenesis and spliceosome disassembly (Wen et al. 2008; Wild et al. 2010). Its specificity is regulated by G-patch co-factors (Heininger et al. 2016), with NKRF controlling rRNA biogenesis (Memet et al. 2017) and TFIP11 mediating spliceosome disassembly (Studer et al. 2020). In this manuscript, we define a new role for DHX15 in early spliceosome assembly that may provide higher eukaryotes an additional quality control mechanism to ensure splicing fidelity in the face of divergent branch sequences. In recent years links between recognition of the branch point region and cancer have dramatically increased (Agrawal et al. 2018). For example, hematological malignancies frequently select for specific point mutations in the U2 snRNP-associated proteins SF3B1 and U2AF1 (Hautin et al. 2020). Notably, DHX15 is also often misregulated in blood cancers (Jing et al. 2018; Pan et al. 2017).

Our working model is that DHX15 performs quality control of the initial interaction between U2 snRNP and an intron by disassembling complexes that are incompetent to form a productive branch helix (Fig 7). It could work in concert with DDX46, which is proposed to directly proofread the branch helix. The yeast ortholog Prp5 releases from A-complex less readily with a mutant vs. consensus branch helix, with its release being necessary for continued spliceosome assembly (Liang and Cheng 2015). If we extend this model to mammalian introns, which have more variable branch point sequences, DDX46 is more likely to stall on nonproductive branch helices. As a result, the more complex intronic landscape necessitates a mechanism for disassembly. We suspect that DHX15 fulfills this role through a disassembly mechanism that parallels disruption of U2 snRNA interactions with the intron in the ILS.

**Figure 7.**
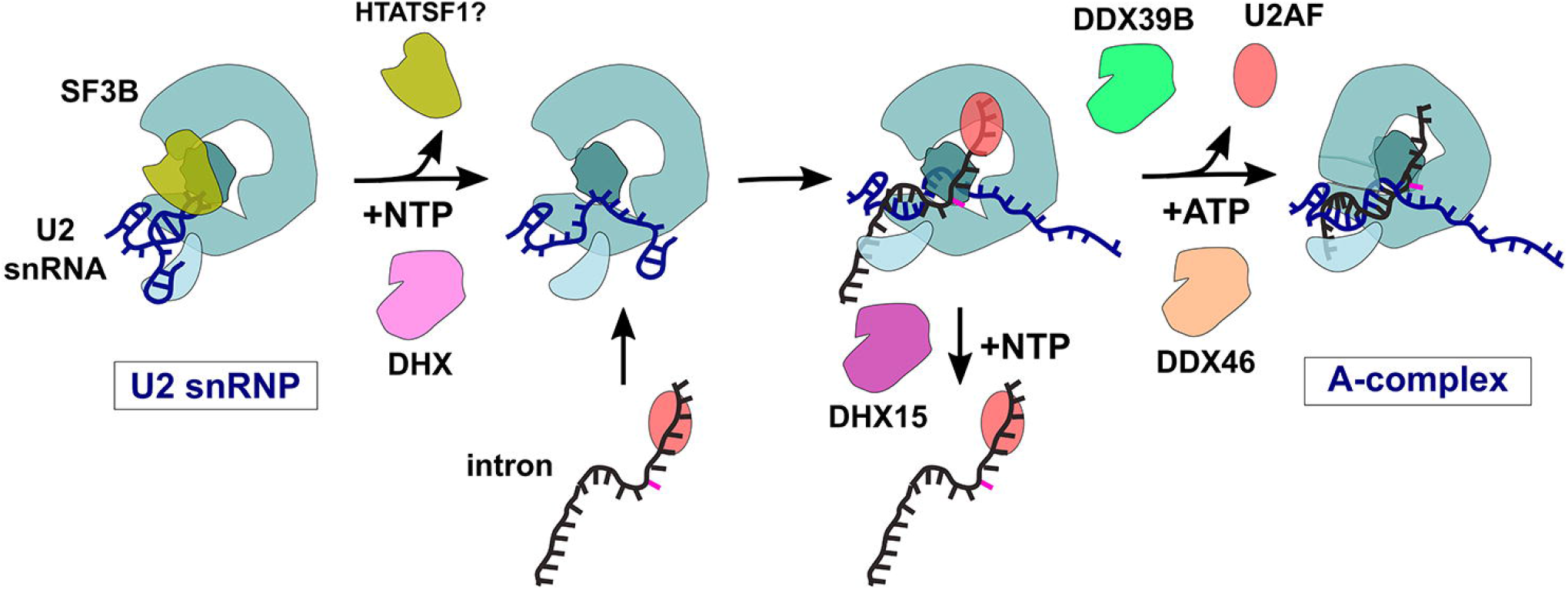
Model of DDX- and DHX-enzyme contributions to early spliceosome assembly.

If DHX15 destabilizes stalled A-complex, how it is recognized comes into question. So far, DHX15’s RNA targets are imposed by the identity of the complex being remodeled. For example, the enzyme promotes ILS disassembly by binding the 3’ end of U6 snRNA (Toroney et al. 2019), although how the branch helix between U2 snRNA and the intron is disrupted remains unclear. A constitutively active DHX15 can also interact with the intron to promote disassembly of B^act^ spliceosomes (Fourmann et al. 2016). In the context the early spliceosome, DHX15’s target is not yet defined. One option is the intron. With the limited real estate of the minimal intron substrate, we were able to narrow down the potential RNA handle to 10 nucleotides downstream from the branch point sequence. U2 snRNA is the other possibility, and we find that the branch binding region of U2 snRNA becomes accessible upon addition of both ATP and GTP. Because DHX15 associates with U2 snRNP, it is a good candidate for mediating the change, which could mimic removal of an intron in either the context of the ILS or A-complex.

Our model also requires a G-patch partner to direct DHX15 to stalled spliceosomes and may further dictate its RNA target. Notably, five different G-patch proteins associate with U2 snRNP and/or A-complex: RBM5, RBM10, RBM17, CHERP and SUGP1. Of these, SUGP1 binds to SF3B1 to prevent use of cryptic branch points, although its DHX-enzyme partner was not identified (Zhang et al. 2019). Mutations in SUGP1 are also associated with aberrant splicing related to branch point selection in cancer cells that also correlate with SF3B1 mutations (Liu et al. 2020).

## Materials and Methods

### HeLa nuclear extract

Nuclear extract was generated as previously described (Dignam et al. 1983) from HeLa S3 cells grown in DMEM/F-12 1:1 and 5% (v/v) newborn calf serum. HeLa cells containing an integrated transgene for V5-tagged SNRPB2 were generated using the HILO-RMCE cassette and RMCE acceptor cells (generously provided by E. Makeyev) as described in Khandelia, *et al*. (Khandelia et al. 2011). Expression of V5-tagged SNRPB2 was induced with doxycycline 48 hours prior to harvest (Kim 2019). Prior to use for the NTP studies, HeLa nuclear extract was incubated at 30°C for 15 minutes to deplete residual ATP.

### T7 run-off RNA transcription

Full-length pre-mRNA sequence is derived from the adenovirus major late (AdML) transcript with uracil substituted at the last four bases of the 5’ exon, and a UACUAAC branch point sequence. Sequences for the A^3’^ and A^min^ RNA substrates are provided in Supplemental Fig. 3. Double stranded DNA oligos (Eurofins) where generated as templates for PYT truncations. All RNA substrates were generated by T7 run-off transcription reactions supplemented with ^32^P-UTP. Full-length substrate was capped by G(5’)ppp(5’)G and gel purified. A^3’^, A^min^ and PYT truncation substrates were GMP-capped and purified by size-exclusion spin column.

### Native gel analysis of *in vitro* splicing reactions

Splicing reactions containing 20 nM RNA substrate, 40 mM potassium glutamate, 2 mM magnesium acetate, 0.05 mg/mL tRNA, 40% (v/v) HeLa nuclear extract and 2 mM ATP, GTP, AMP-PNP or no added NTP were incubated at 30°C for 0-15 minutes. 2X native loading dye (20mM Tris, 20mM glycine, 25% glycerol, 0.1% bromophenol blue, 0.1% cyan blue, 1mg/ml heparin) was added directly to each reaction, which were separated on native gels with 1.2% low melting temp agarose (Invitrogen, 15517-014) in 20 mM Tris, 20 mM glycine for 3 hours at 72 volts. Gels were dried onto Whatman paper and visualized by phosphorimaging (Amershan Typhoon).

### Denaturing gel analysis of *in vitro* splicing reactions

Splicing reactions were generated as described above except that pre-mRNA substrate was at 6 nM and incubation time was extended to 45 minutes. Reactions were stopped with addition of splicing dilution buffer (100 mM Tris pH 7.5, 10 mM EDTA, 1% SDS, 150 mM sodium chloride, and 0.3 M sodium acetate pH 5.2) followed by phenol:chloroform:isoamyl alcohol (25:24:1) extraction and ethanol precipitation. RNA was resuspended in FEB (95% formamide, 20 mM EDTA, 0.01% bromophenol blue and 0.01% cyan blue), separated by 15% urea-PAGE and visualized by phosphorimaging.

### DHX15 depletion

10 µg DHX15 antibody (Abcam, ab70454), IgG (Epredia™, NC748P) or 1X PBS was incubated with protein A beads (NEB, S1425S) with agitation for 1 hour at 4°C. HeLa nuclear extract with 500 mM potassium chloride was added to beads and rotated end over end for 2 hours at 4°C. The depleted nuclear extract was then dialyzed into Buffer E for 5 hours (100 mM potassium chloride, 20% glycerol, 20 mM Tris, pH 7.9, 1.5 mM magnesium chloride, 0.5 mM DTT) in small dialysis cups (Thermo Scientific™, 69570).

### RNase H digestion and primer extension

HeLa nuclear extract was supplemented with 2 mM magnesium acetate and 2 mM ATP, GTP, AMP-PNP or no NTP. DNA oligos (Eurofins) complementary to the U2 snRNA nt 1-15, nt 12-26, nt 24-38 and nt 32-46 or a control oligo was added at 5 µM and incubated for 15 minutes at 30°C to allow for cleavage by endogenous Rnase H. The oligonucleotides were degraded by addition of 1µl RQ1 Dnase (Promega, M6101) for 10 minutes at 30°C. RNA was isolated by phenol:chloroform:isoamyl alcohol (25:24:1) and chloroform:isoamyl alcohol (24:1) extraction, followed by ethanol precipitation and resuspended in water. For primer extension, 10 picomoles of DNA oligonucleotides complementary to either U2 snRNA nt 97-117 or nt 28-42 (U2L15 (Black et al. 1985)) were labeled with γ-32P ATP and purified via Sephadex G-25 (MilliporeSigma™ Supelco™ G258010G) column. The isolated RNA and radiolabeled oligonucleotide were annealed by incubation at 95°C for 2 min, 53°C for 5 min, and on ice for 5 minutes and then added to reverse transcription reactions containing 50 mM Tris pH 7.9, 75 mM potassium chloride, 7 mM DTT, 3 mM magnesium chloride, 1 mM dNTPs, and 0.5 µg reverse transcriptase (MMLV variant). Reactions were incubated for 30 minutes at 53°C, and the DNA was isolated by the addition of 0.3 M sodium acetate pH 5.2, 0.5 mM EDTA, and 0.05% SDS followed by ethanol precipitation. The labeled DNA was resuspended in FEB and separated on a 9.6% urea:PAGE. The gel was dried on Whatman paper and visualized by phosphorimaging.

### Immunoprecipitations

For IP’s in Fig 6A, C & D, 1.5 µg of anti-V5 antibody (Invitrogen, R960-25) or IgG (Epredia™, NC748P) was added directly to splicing reactions containing 6 nM full-length RNA substrate, 2 mM magnesium acetate, 40 mM potassium glutamate, 0.1 mg/ml tRNA and 40% V5-SNRPB2 HeLa +/-2 mM ATP incubated at 30°C for 10 minutes. For other co-IP’s, 3 µg of anti-V5 antibody (Genescript, A01724) was added to 2 mM magnesium acetate, 20 mM potassium glutamate, 0.1 mg/ml tRNA and 60% V5-SNRPB2 HeLa nuclear extract +/– 2 mM ATP, GTP or AMP-PNP and incubated at 30°C for 10 minutes. All samples were incubated with antibody for 13.5 hours at 4°C with rocking. Samples were then added to protein A magnetic beads (NEB, S1425S) and rotated at 4°C for 4 hours. The beads were washed three or more times with IP wash buffer (100 mM Tris pH 7.5, 120 mM potassium chloride, 1 mM EGTA, 0.1% NP-40 or IGEPAL). For western blots, samples were eluted with 0.1 M glycine, pH 2.5 and quenched with equal volume 1 M Tris, pH 7.9. For RNA analysis, samples were eluted with splicing dilution buffer followed by precipitation with phenol:chloroform:isoamyl alcohol (25:24:1) and ethanol precipitation. RNA samples were analyzed by 15% urea-PAGE as described for *in vitro* splicing reactions.

### Western blot analysis

Samples were prepared in 5X Laemmli buffer (62.5 mM Tris, 25% glycerol, 6.25% SDS, 0.1% bromophenol blue, 5% beta-mercaptoethanol) and heated at 95°C for 2 minutes prior to separation by 10% SDS-PAGE. Gels were transferred to PVDF membrane (Bio-Rad Mini Trans-Blot® Cell) and blocked in 1% non-fat milk in 1X Tris-buffered saline with Tween 20 (TBST) for 1 hour at room temperature while rocking. The following antibodies were added directly to the blocking buffer at the indicated concentrations and incubated at 4°C overnight while rocking. From Proteintech: DHX15 (12265-1-AP, 1:1000), SNRPB2 (13512-1-AP, 1:2500), U2AF1 (60289-1-Ig, 1:1000). From Santa Cruz Biotech: SF3A2 (sc-390444, 1:1000). Blots were washed three times in 1X TBST and corresponding LICOR secondaries (1:15000) were added in blocking buffer and rocked at room temperature for 1 hour. The blots were again washed three times and then imaged on a LICOR Imaging System. Images were quantified and processed utilizing LICOR Image Studio Lite.

## Acknowledgements

This work was supported by a National Institutes of Health grant R01GM72649 to M.S.J.

## Author Contributions

Conceived and designed the experiments: HMN, AA, JK, MM, MJ. Performed the experiments: HMN, AA, TS, JK, MM, BP. Analyzed the data: HMN, AA, JK, MJ. Wrote the paper: HMN, AA, MJ.

## Competing interests

The authors have declared that no competing interests exist.

## Notes

### Competing Interest Statement

The authors have declared no competing interest.

